# Prostate-specific EV capture with sufficient RNA yield to enable transcriptomic profiling

**DOI:** 10.1101/2025.09.08.674937

**Authors:** Yang Yang, Julia Doo, Devin Perez, Kurt Franzen, Sinead Nguyen, Jasmyn Mirabal, Emily Mistock, Sanoj Punnen, Sandra Gaston, Timothy Gerard, Alan Pollack, Elena Cortizas, Sudipto K. Chakrabortty, Johan K. Skog, Joseph M. Johnson

## Abstract

Prostate-specific antigen (PSA) screening has reduced prostate cancer (PCa) mortality but suffers from limited specificity, contributing to unnecessary biopsies and overdiagnosis of indolent disease. There is a critical need for biofluid-based biomarkers that improve the precision of PCa detection. Extracellular vesicles (EVs) offer a promising platform for noninvasive diagnostics, as they carry molecular cargo reflective of their tissue of origin. The ExoDx Prostate IntelliScore (EPI) test, a urine-based EV assay, is currently the only commercial EV diagnostic for clinically significant (cs)PCa, but its performance may be constrained by contamination from renal and bladder-derived EVs.

To address this, we developed Exosome Diagnostics Depletion and Enrichment (EDDE), a novel immunocapture-based method for isolating prostate-derived EVs with high specificity. By targeting Prostate Specific Membrane Antigen (PSMA), we optimized EDDE to selectively enrich prostate EVs from post-DRE urine and recover sufficient RNA for transcriptomic analysis. Throughout development, we implemented a quantitative framework to track EV stoichiometry and assess depletion efficiency and yield, enabling rigorous optimization of the workflow.

Our findings demonstrate that PSMA EDDE enriches prostate-specific EVs and yields RNA quantities compatible with sequencing. This platform enhances the specificity of EV-based biomarker discovery and holds promise for determining if tissue-specific EV biomarkers contribute advantages over bulk EVs.

## Introduction

Prostate-specific antigen (PSA) screening reduces prostate cancer (PCa) mortality but lacks specificity, leading to unnecessary biopsies and overdiagnosis of indolent disease (Loeb et al., 2011; Schröder et al., 2012). Multi-parametric MRI (mpMRI) improves detection of clinically significant PCa (csPCa), yet its interpretation remains subjective and prone to false positives and negatives (Ahdoot et al., 2020; Eklund et al., 2021; Kasivisvanathan et al., 2018; Westphalen et al., 2020). Commercially available urine and blood-based biomarkers aim to refine biopsy decisions by estimating the probability of csPCa, typically defined as Grade Group 2 or higher (Becerra et al., 2019, 2021). However, limited specificity continues to drive excessive biopsy rates (Falagario et al., 2021). There is a *critical need* for biofluid based biomarkers that can substantially enhance the specificity of PCa evaluation and reduce the burdens of PCa screening.

Extracellular vesicles (EVs) offer a promising platform for noninvasive cancer diagnostics. Tumor cells release EVs into biofluids, packaging molecular cargo—including RNA and proteins—within lipid bilayers that preserve their integrity (Yu et al., 2021). The EPI test from Exosome Diagnostics, the only FDA-approved EV-based PCa diagnostic, uses qRT-PCR to measure three RNA biomarkers in urine-derived EVs to assess csPCa risk (Loeb et al., 2011). Urine has reduced contamination from non-genitourinary EVs, but still contains EVs from renal and bladder sources, which may dilute prostate-specific signals. We hypothesize that analyzing prostate-specific EVs will improve csPCa biomarker performance compared to total urine EVs (Kim et al., 2022; Lucien et al., 2022).

Several studies have claimed successful isolation of tissue-specific EVs, yet many of these reports have faced significant scrutiny. Multiple publications describing L1CAM-based capture of neuron-derived EVs in plasma and cerebrospinal fluid were later challenged by rigorous analytical work showing that L1CAM is not associated with EVs in these fluids (Gomes & Witwer, 2022; Kapogiannis et al., 2019; Norman et al., 2021; Yan et al., 2023). Similarly, reports of detecting pathological α-synuclein aggregates in blood-derived neuron-specific EVs have been called into question due to the use of antibodies that target the cytoplasmic protein domain of L1CAM (Kluge et al., 2022; Bernhardt et al., 2024). These controversies highlight the pitfalls of asserting tissue-specific EV isolation without thorough analytical validation. In the context of prostate cancer, PSMA-based EV enrichment has shown promise, but optimal conditions for EV capture and processing remain undefined, and prior protocols have not undergone robust diagnostic development (Allelein et al., 2022; Wang et al., 2023).

To test our hypothesis, we developed Exosome Diagnostics Depletion and Enrichment (EDDE), a workflow that enables transcriptomic analysis of immuno-captured prostate EVs from urine (Figure 1). To ensure reliable tissue-specific EV isolation, EDDE incorporated a multi-tiered development process including screening of multiple surface targets before choosing prostate specific membrane antigen (PSMA), establishing sample and process controls, screening bead types and optimization of surface chemistry for low non-specific binding (nsb), screening for optimal capture antibodies, and optimization of buffer conditions. Perhaps most importantly, to evaluate EDDE performance we have established a framework for quantitatively tracking EV stoichiometry during enrichment and quantitatively measuring both protein and RNA yield.

**Figure 1.**
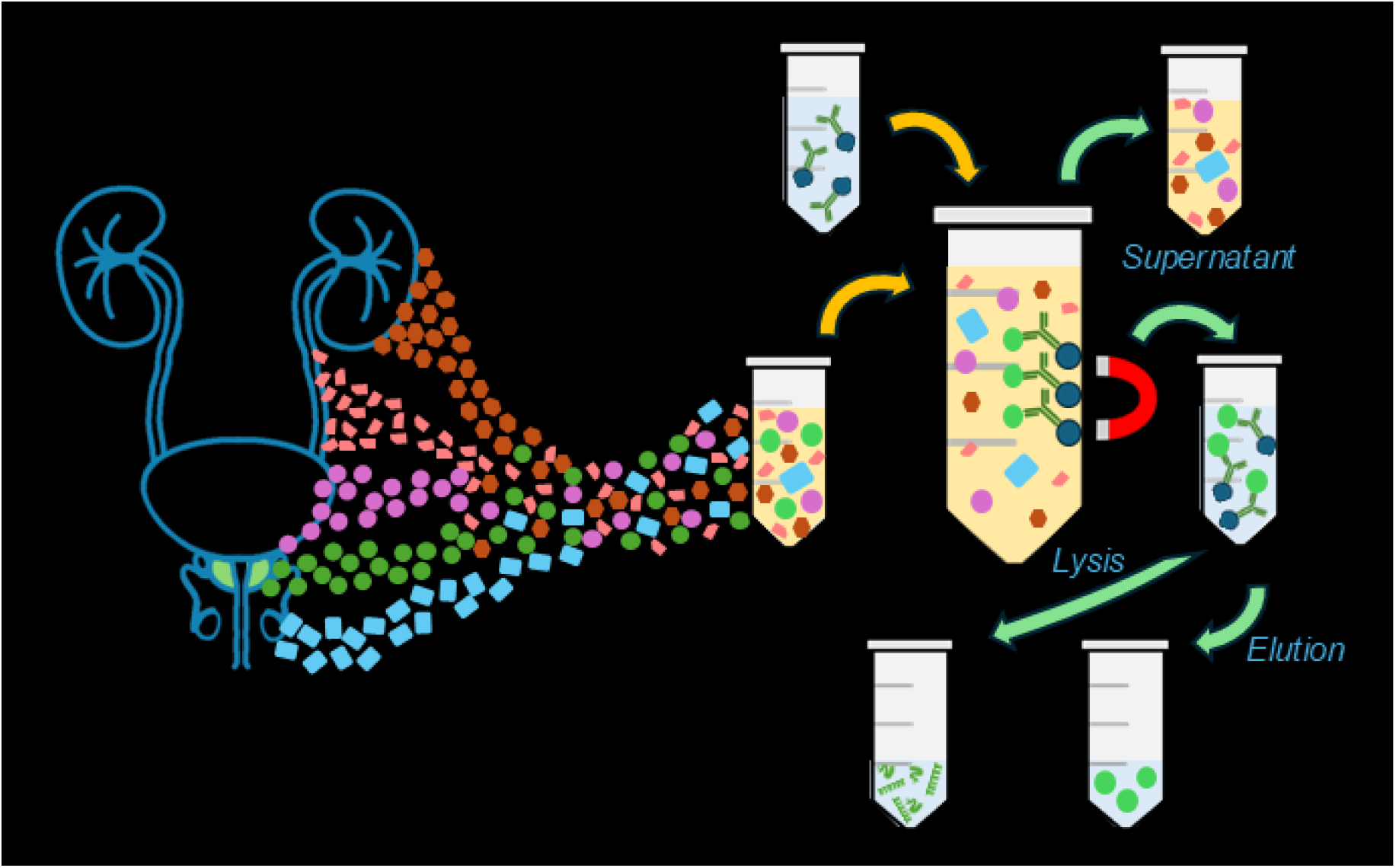
EDDE captures tissue-specific EVs. Mixtures of EVs can be separated by EDDE, with downstream analysis of either the supernatant, intact EVs (Elution) or cargo molecules (Lysis).

We applied EDDE to samples from the ongoing MDSelect trial (NCT04240327), which enrolls men undergoing biopsy for csPCa evaluation. Our optimized protocol yields sufficient RNA from prostate-specific EVs to support transcriptomic profiling using single-cell RNA sequencing workflows. This study represents the first demonstration of prostate-specific EV capture with sufficient yield to enable RNA sequencing and lays the foundation for biomarker discovery with enhanced tissue specificity.

## Materials and Methods

### Clinical sample collection

The MDSelect trial (NCT04240327) is an ongoing NCI funded prospective protocol enrolling men referred for biopsy of their prostate for evaluation of PCa at U Miami. All men undergo mpMRI, interpreted with standard of care PIRADS, prior to biopsy of the prostate. Patients provide blood and urine samples prior to biopsy, which are sent prospectively for a 4KScore and EPI test. All patients undergo systematic template sampling of the prostate (12 cores) during biopsy in addition to targeted biopsy of any mpMRI suspicious regions. MDSelect will enroll 250 men who will undergo MRI guided biopsy for prostate cancer detection. The trial is funded to evaluate a quantitative MRI imaging algorithm and will result in the generation of key MRI features and radiomic variables related to the presence of csPCa. The primary outcome of the protocol is the detection of csPCa, defined as GG2+ cancer on biopsy of the prostate. These men will serve as intermediate to high-risk cases in Specific Aims 2 and 3 of this proposal. Men with no cancer or indolent cancer (GG 1) will be used as low-risk controls. Based on an expected GG2+ detection rate of 40%, we expect 100 men with GG2+ cancer (cases) and 150 men with indolent or no cancer (controls). The MDSelect trial is meeting enrollment, and among the 64 men who have undergone biopsy to date, 33 (52%) had csPCa, and 31 (48%) had indolent or no cancer.

### Materials

Antibodies ExoView staining and EDDE: anti-PSMA (BioLegend clone LNI-17 catalog# 342502; R&D Systems clone 460407 catalog# MAB42342 and FAB42342R; R&D Systems polyclonal catalog# AF4234; Millipore Sigma clone 3/A12 catalog# MABC291); anti-SLC45A3 (Abcam clone EPR4795(2) catalog# ab318068; Santa Cruz Biotechnology clone A-5 catalog# sc-393069 PE); anti-KLK2 (LSBio monoclonal catalog# LS-C51453; Novus Biologicals clone JNJ-69086420 catalog# NBP3-28696AF647); anti-CD81 (R&D Systems clone 454720 catalog# MAB4615); isotype control mouse IgG (R&D Systems clone 11711 catalog# MAB002). R&D Systems polyclonal anti-PSMA and Millipore Sigma anti-PSMA were labeled with Alexa Fluor 555 (Thermo Fisher, catalog# A20187) and Alexa Fluor 647 (Thermo Fisher, catalog# A20186).

Purified LNCaP EVs (ATCC, catalog# CRL-1740-EXM) and PC3 EVs (ATCC, catalog# CRL-1435-EXM) were used as PSMA+ and PSMA-controls, respectively.

### Beads and antibody conjugation

EDDE beads were prepared using a proprietary method.

### EDDE with LNCaP EV and Urine

EDDE was performed with a proprietary method. Briefly, for EDDE with urine samples, 100 µL of the antibody conjugated beads were mixed with 5 or 7 mL of the urine samples. Subsequently, the standard EDDE procedure was followed for washing and lysis. In addition, 200 µL of urine samples and EDDE supernatants were lysed using the QIAzol Lysis Reagent to assess RNA in the pre-EDDE samples and determine the remaining RNA in the supernatants after the EDDE process.

For serial EDDE, the supernatants were mixed again with 100 µL of the antibody conjugated beads. The EVs were then eluted or lysed following the same procedure as described above. Three consecutive EDDE runs were conducted to evaluate the performance of EV enrichment.

### ExoView (rebranded Leprechaun)

Samples were diluted in PBS containing 0.5% BSA with or without 0.1% Tween-20 and sample diluent provided in the Leprechaun Human Tetraspanin Kit (Unchained Labs, catalog# 251-1044). A higher sample dilution (e.g., 10,000 to 25,000-fold for purified EVs and 6 to 320-fold for urine samples) was necessary for the measurement of total tetraspanins (i.e., CD9, CD63, and CD81), while a lower dilution (e.g., 50 to100-fold for purified EVs and 2-fold for urine samples) was required for specific markers such as PSMA, SLC45A3, and KLK2.

Fifty microliters of the samples were then added to Leprechaun chips and incubated overnight at room temperature. The chips were subjected to automated washing and fluorescent staining using the ExoView chip washer (NanoView, catalog# CW100). Subsequently, the chips were scanned using the ExoView chip scanner (NanoView, catalog# R200) and the data were analyzed using the ExoView Analyzer software (version 3.2.1). The limit of detection for each assay was taken as the EV count on the MIgG isotype control capture probe, multiplied by two.

During data analysis, a predetermined cutoff value was established for each fluorescent channel and maintained consistently across all samples when measuring the same target. The cutoff value typically fell within the range of 300-400 for the AF647 and AF555 channels, and between 500-600 for the AF488 channel. Only EVs exhibiting fluorescent signals surpassing the respective cutoff values were included in the counts.

### On-bead enzyme-linked immunosorbent assay (ELISA)

As part of a proof-of-concept experiment, EDDE was performed with slight modifications to establish an on-bead ELISA. Twenty microliters of the antibody conjugated beads were added to individual tubes, followed by the removal of supernatants. Subsequently, a serial dilution of PSMA calibrator from the PSMA kit (Meso Scale Discovery, catalog# K151ABPR) ranging from 100 ng/mL to 0.1 ng/mL was prepared. The samples were then incubated with 0.1 mL of PSMA standards and 50-fold diluted LNCaP EVs, respectively, for 1 hour at room temperature with rotation. A 50-fold diluted pre-lysed LNCaP EVs was also included to determine total PSMA concentration in the sample. After incubation, the beads were resuspended in 0.1 mL of PBS and transferred to a fresh tube. The beads were washed twice with wash buffer (0.1 mL of 0.1% Tween-20 in TBS). Following this step, the EVs attached to the beads were lysed by adding 20 µL of 0.2% Triton X-100 (Sigma-Aldrich, catalog# X100-5ML), vortexing for 30 seconds, and incubating for 15 minutes at room temperature with rotation. The beads were washed twice again. For the subsequent on-bead ELISA, 100 µL of 1 µg/mL biotinylated polyclonal anti-PSMA (R&D Systems, catalog# BAF4234) was added to the beads and incubated for 1 hour at room temperature with rotation. The beads were washed twice, and streptavidin-horseradish peroxidase (HRP, R&D Systems, catalog# DY998) was diluted to the working concentration specified on the vial label using 1X Reagent Diluent Concentrate 2 from DuoSet Ancillary Reagent Kit 2 (R&D Systems, catalog# DY008B). Subsequently, 20 µL of the streptavidin-HRP working solution was added to the beads and incubated for at least 20 minutes at room temperature with rotation. After the last washing step of the beads, the SuperSignal ELISA Femto Luminol/Enhancer and SuperSignal ELISA Femto Stable Peroxide Solution (Thermo Scientific, catalog# 37075) were mixed in equal parts. Two hundred microliters of working solution were added to the beads, and the solution containing the beads was transferred to a 96-well plate for a 1-minute incubation. Relative light units were measured at 425 nm between 1-5 minutes after adding the working solution.

### Meso Scale Discovery Electrochemiluminescence Immunoassay

Meso Scale Discovery Electrochemiluminescence Immunoassay (MSD) was performed for measurement of intact CD81 EV concentration with R-PLEX Human CD81 (EV) Assay (Meso Scale Discovery, catalog# K1515NR-2) and PSMA protein concentration with R-PLEX Human GCPII/PSMA Assay (Meso Scale Discovery, catalog# K151ABPR-2). MSD assays were performed according to the manufacturer’s instructions with the exceptions noted below. Capture antibody was coated on a 96-well plate, followed by overnight sample incubation with shaking at 4 °C. For CD81 EV measurement with R-PLEX Human CD81 (EV) Assay, samples and calibrators were diluted with Diluent 52 (Meso Scale Discovery, catalog# R52AA-1) with addition of 0.5 % BSA and 0.1% Tween-20. LNCaP EVs were used as calibrators for the CD81 EV assay, with a value assignment of 1 × 10^12^ EV/mL stock concentration, as measured by ExoView platform with Leprechaun Human Tetraspanin Kit (with a range of 3.4 × 10^11^ to 4.0 × 10^12^ EV/mL over four experiments (Supplemental Figure 1). For PSMA protein measurement with R-PLEX Human GCPII/PSMA Assay, samples and calibrators were diluted in Diluent 11 (Meso Scale Discovery, catalog# R55BA-5) with addition of 0.5 % Triton X-100 (Sigma-Aldrich, catalog# X100-5ML). This diluent lysed EVs and reduced the assay background, enabling a lowered limit of detection (LoD) than what was claimed by the manufacturer. MSD plates were measured on a MESO QuickPlex SQ 120MM (Meso Scale Discovery, catalog# AI1AA-0) and analyzed MSD Discovery Workbench software (version 4.0.13). Standard curves were generated using Graphpad Prism (version 10.6.0) by fitting data with four-parameter logistic regression and applying 1/Y^2^ weighting. Subsequently, the concentrations of unknown samples were backcalculated based on the established standard curves.

### RNA quantification

Five to ten mL of urine were filtered with 0.8 µm Millex AA filter units (Millipore Sigma) and an exogenous RNA extraction control was spiked in. EV isolation and RNA extraction was performed using ExosomeDx’s proprietary ExoLution RNA workflow. Total RNA isolated from EVs was reverse transcribed using Superscript IV VILO MasterMix (Thermo Fisher Scientific). Reverse transcription was carried out under thermocycle conditions optimized for conversion of EV-derived RNA fragments to cDNA [25 °C for 10 min, 48 °C for 30 min, 85 °C for 5 min, followed by a 4 °C hold]. cDNA was used immediately or stored at 4 °C overnight.

Quantitative PCR (qPCR) was performed using cDNA generated from the reverse transcribed EV-isolated RNA, TaqMan Fact Universal PCR Master Mix: 2x No AMPErase UNG (Thermo Fisher Scientific), and either 20x TaqMan Gene expression qPCR Assays (Thermo Fisher Scientific or IDT) or proprietary custom qPCR Primers and Probes (ExosomeDx; synthesized by IDT). Commercial assays used included: FOLH1 Hs00379515_m1 (Thermo Fisher), RPL11 Hs.PT.58.15587787, RPSA-1 Hs.PT.58.3211766, RPL27 Hs.PT.58.28276759, and RPSA18S Hs.PT.58.2672875 (IDT). PCR was performed on QuantStudio 5 RealTime PCR systems [95 °C for 20 seconds, 40 cycles of 95 °C for 1 second and 65 °C for 20 seconds]. Amplification data were analyzed using QuantStudio Design & Analysis Software (Thermo Fisher Scientific). A threshold setting of 0.1 and auto baseline correction were applied uniformly across all runs.

For quantification, RNA isolated from urine-derived EVs was quantified using an Agilent Bioanalyzer with the RNA Pico 6000 Kit (Agilent Technologies, catalog# 5067-1513) in duplicate over two runs (Supplemental Figure 2). Multiple aliquots of the RNA stock were stored frozen, and standard curves prepared from them were included for each assay in every run. Measured Ct values plotted against RNA concentrations were fit with a four-parameter logistic regression, which was used to convert Ct values of unknowns to RNA mass in picograms.

## Results

### Selection of a surface antigen for prostate-specific EVs

For initial EV characterization in samples, we used the ExoView system which captures EVs with anti-tetraspanin antibodies and visualizes them after staining for surface proteins (Figure 2). The ExoView system also incorporates sizing information via interferometry measurements, ensuring that the measured samples are <200 nm in diameter. Measurements of EVs on ExoView therefore fulfill many of the MISEV requirements directly, as it only measures tetraspanin+ particles within the correct size region for EVs (Welsh et al., 2024).

**Figure 2.**
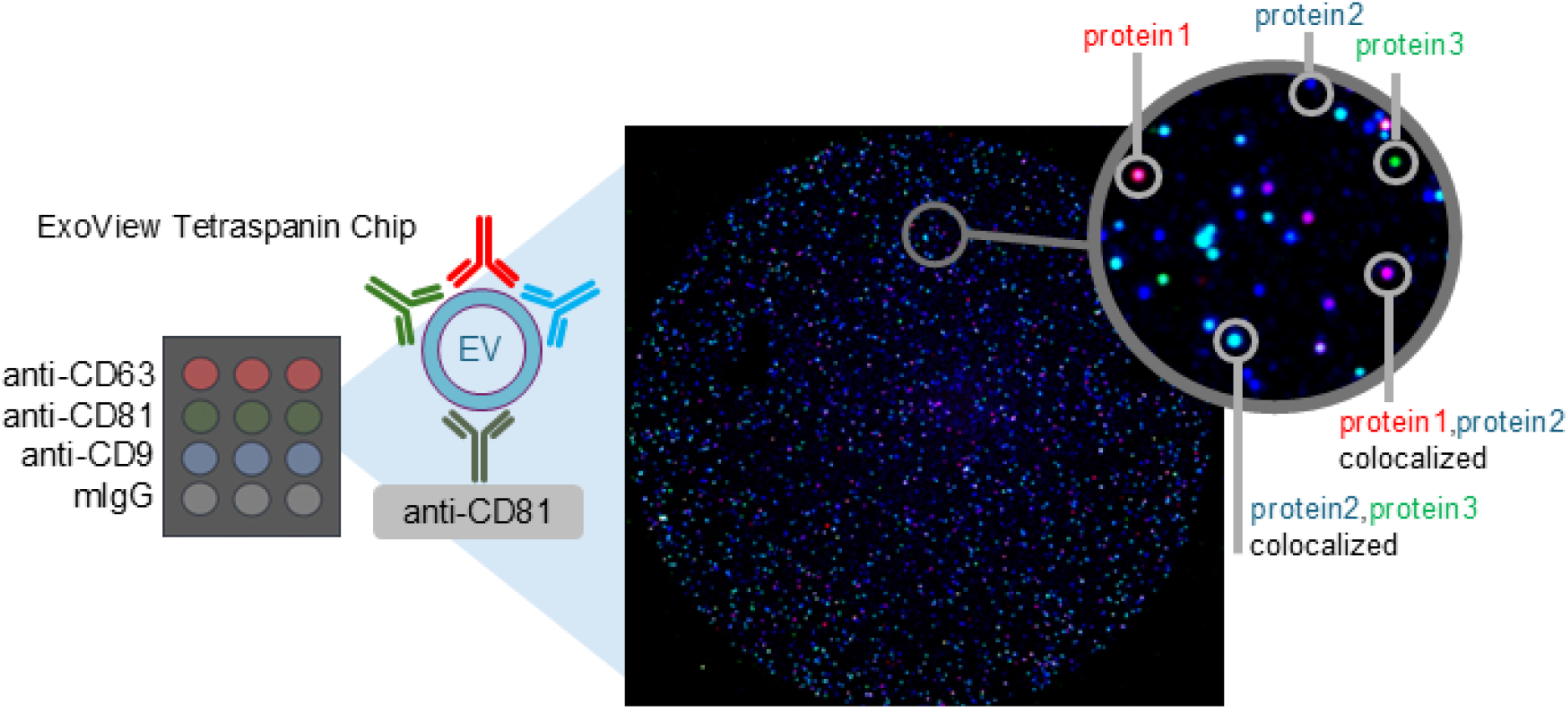
ExoView analyzes EVs according to MISEV guidelines. A schematic of the ExoView capture chip, an actual image and zoomed close-up showing individual stained particles. The color corresponds to the protein stained.

To select the optimal target for prostate EV enrichment, we surveyed urine samples from PCa patients on ExoView. We quantified total EV concentrations in urine samples from men with Grade Group 0 (GG0, n=10) and Grade Group 5 (GG5, n=10) using ExoView analysis (Figure 3**Error! Reference source not found**.A). We found that CD81+ EVs are lower in urine samples compared to CD63+ or CD9+ EVs, and then total EVs are lower on average in GG5 than GG0 urine. We also quantitated prostate-specific EV concentration by measuring PSMA+ (n=20), SLC45A3+ (n=20) and KLK2+ EVs(n=4) (Figure 3**Error! Reference source not found**.B). We found that PSMA+ EVs and SLC45A3+ EVs had the highest concentration relative to total EV, both making up 0.05% of the total EV concentration on average. However, the SLC45A3 assay had a higher LoD. as determined by non-specific binding to the MIgG capture probe and many of the individual samples fell below this. From these results, we selected PSMA as the surface target for enriching prostate-specific EVs. At least two antibody clones were evaluated for each target before screening samples (Materials).

**Figure 3.**
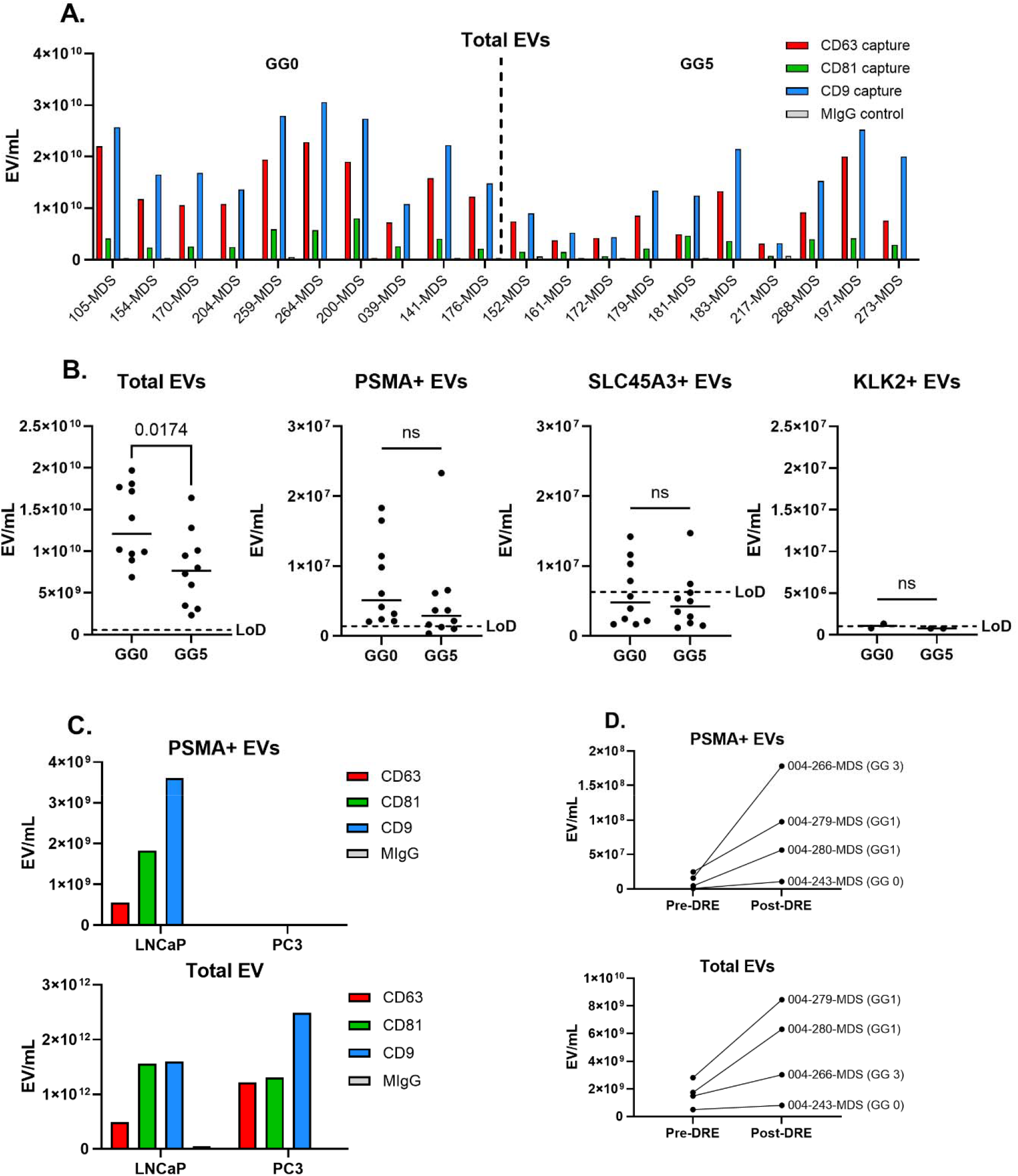
PSMA is the most promising surface target surveyed for prostate EV enrichment. Panel A – Total EV concentration in PCa urine samples analyzed with ExoView for GG0 (n=10) and GG5 (n=10) donors. Concentrations were measured with CD63, CD81 and CD9 capture probes, and detected with a cocktail of detection antibodies to the same three tetraspanins. Panel B – PCa urine samples measured on ExoView for EVs containing multiple surface markers including total EVs (n=20), PSMA+ EVs (n=20), SLC45A3+ EVs (n=20), and KLK2+ EVs (n=4). The data is averaged over all three capture probes (CD63, CD81, CD9). The LoD for each assay is shown as a dashed line. Panel C – PSMA+ and total EV concentrations measured in purified EVs from LNCaP or PC3 cell lines. Panel D. – PSMA+ and total EV concentrations measured in urine samples from PCa donors before (n=4) and after prostate massage (n=4); average of capture probes is shown.

LNCaP and PC3 are known to be PSMA+ and PSMA-PCa cell lines respectively (Ghosh et al., 2005), and we verified this using total and PSMA+ EV Exoview measurements (Figure 3**Error! Reference source not found**.C). We made use of purified EVs from these cell lines available from ATCC as a positive and negative control for development of a PSMA pull down process. Even for the known positive sample LNCaP, PSMA+ EVs made up only a small portion of the total EV population (<1%).

We also measured total and PSMA+ EVs from pre- and post-DRE (digital rectal exam) urine from PCa donors (Figure 3**Error! Reference source not found**.D, n=4). We found an average increase of 2.6-fold for total EVs and 11.7-fold for PSMA+ EVs post-DRE, making this a promising approach for increasing the amount of PSMA+ EVs available for enrichment.

### Optimization of target-specific EV isolation

To facilitate development of EDDE we first evaluated the performance of positive and negative process controls, EDDE CD81 and EDDE IgG, using LNCaP-derived EVs. Figure 4A shows total EV counts measured by ExoView across EDDE fractions: sample pre-enrichment (PreEDDE), supernatant (Sup.) and eluate. EDDE CD81 depleted 94% of EVs from the supernatant and recovered 24% in the eluate. The isotype control EDDE IgG shows minimal (7%) non-specific binding (nsb) and no detectable yield of EVs in the eluate, as expected. Figure 4B confirms these findings using an orthogonal measurement method, MSD-CD81 intact EV detection, in a replicate experiment in which we see similarly efficient CD81 EV capture and low non-specific binding. LNCaP EVs subjected to multiple rounds of EDDE CD81 showed increasingly lower levels of EVs in the supernatant after each round, as expected (Supplemental Figure 3). These experiments validate the robustness of CD81-based EV capture but also indicate inefficient EV elution off the beads.

**Figure 4.**
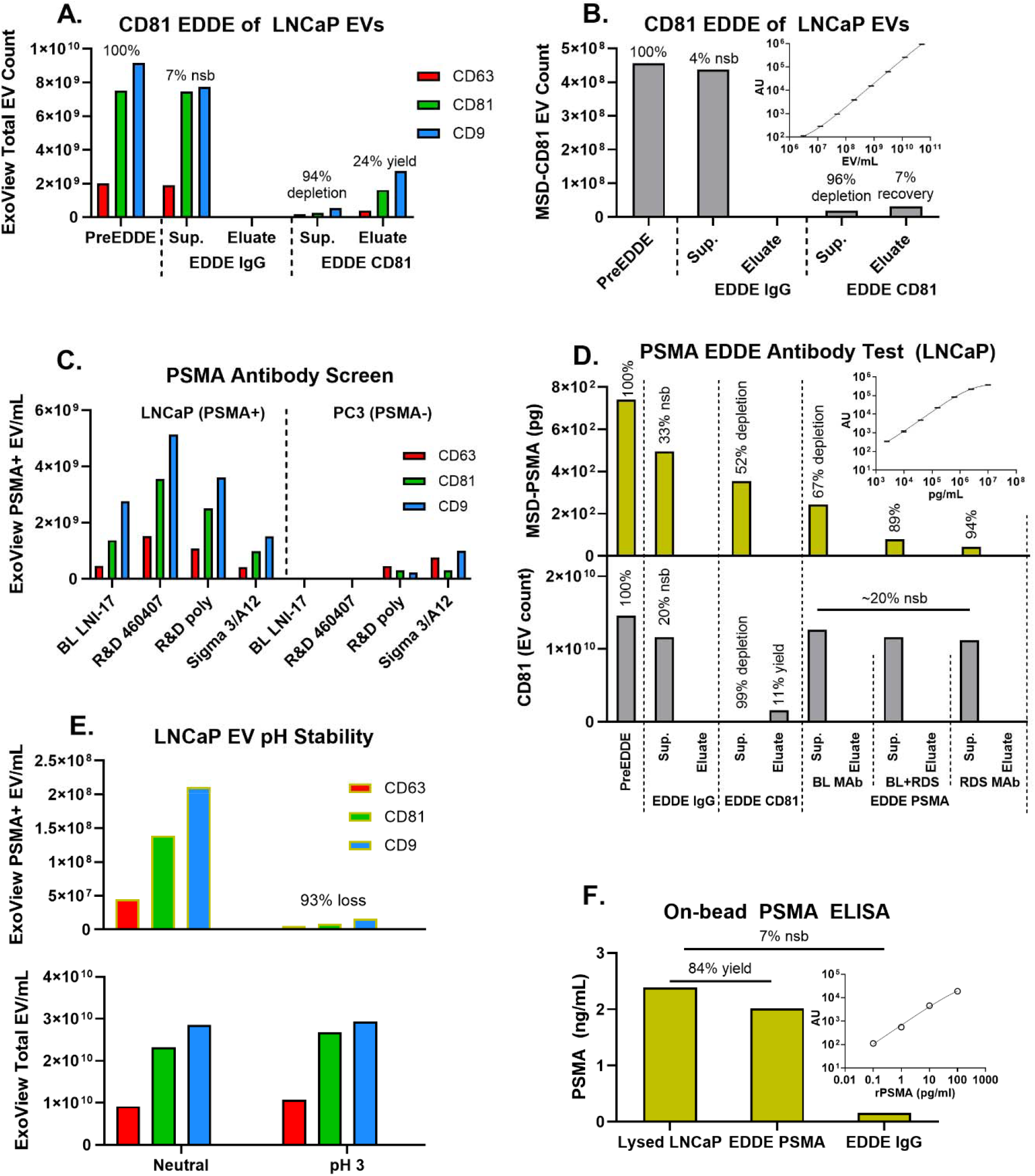
EDDE was optimized using process controls for prostate specific targets. Panel A – EDDE CD81 (+ control) and EDDE IgG (-control) were carried out on purified LNCaP EVs and readout for total EVs on ExoView. The average EV counts from all capture probes (CD63, CD81, CD9) are used to calculate depletion and eluate yield. Panel B – A replicate experiment of EDDE CD81 and EDDE IgG read out with the MSD-CD81 intact EV assay, showing similar levels of depletion and EV yield as ExoView. The insert shows the calibration curve obtained with a dilution of LNCaP EVs at known concentration. Panel C – A screen of PSMA antibodies using LNCaP (PSMA+) and PC3 (PSMA-) purified EVs on ExoView to identify the best antibody. Panel D – Follow-up EDDE test of the two best performing PSMA antibodies and a mixture of the two, as read out with MSD-PSMA and MSD-CD81. The inset shows the calibration curve with recombinant PSMA. Panel E – A stability test demonstrating loss of PSMA+ EV detection at low pH. Total EVs are not affected. Panel F – PSMA on-bead ELISA demonstrating capture of PSMA EVs on beads using EDDE PSMA, with minor non-specific binding observed for isotype IgG beads.

Having established working process controls, we proceeded to identify the most effective antibody for PSMA+ EV enrichment by screening multiple clones using ExoView. We compiled a list of most commercially available antibodies and their known epitopes (Supplemental Figure 4) (Davis et al., 2005). Figure 4C shows PSMA+ EV concentrations in LNCaP (PSMA+) and PC3 (PSMA−) EVs across four antibody candidates. Clones LNI-17 and 460407 demonstrated the highest specificity and signal-to-noise ratio, distinguishing PSMA+ EVs in LNCaP while minimizing background in PC3.

We then evaluated the performance of PSMA EDDE with these clones and a mixture of them (Figure 4D), as read-out with MSD-PSMA (top graph) and MSD-CD81+ EV assay (bottom graph). The clone 460407 had very efficient PSMA depletion from LNCaP EVs (94%). CD81 EDDE showed the expected depletion and eluate yield of CD81+ EVs, while EDDE IgG and all the EDDE PSMA conditions resulted in a moderate amount of non-specific depletion of CD81+ EVs.

We did not detect PSMA in EDDE eluates under any conditions, despite seeing modest yield of CD81+ EVs in eluate. To assess EV stability under acidic conditions used in elution, we compared total and PSMA+ EV concentrations at neutral pH and pH 3. Figure 4E shows minimal effect on total EV concentration but 93% loss of PSMA+ EVs at pH 3, explaining why we failed to detect PSMA+ EVs in any eluate.

To confirm capture of PSMA+ EVs, we developed an on-bead ELISA to measure PSMA protein in lysed EVs after EDDE pulldown with the following workflow: PSMA+ EVs were captured onto EDDE beads which were then washed; captured EVs were lysed with detergent; PSMA was then detected on the bead with (Figure 4F). Compared to the total amount of PSMA signal in the lysed sample, we saw 84% of PSMA retained on-bead after PSMA EDDE, confirming that EDDE both depleted and retained the majority of PSMA+ EVs from LNCaP EVs. We observed a modest 7% non-specific binding signal from EDDE IgG.

### Quantification of RNA from EDDE

We applied PSMA EDDE to post-DRE urine samples which have enhanced levels of PSMA+ EVs (Figure 3**Error! Reference source not found**.D), to selectively enrich prostate-derived EVs and assess their RNA content. The MSD-PSMA immunoassay confirmed reasonable depletion of PSMA protein during the EDDE process, although there is sample-to-sample variability, and virtually no non-specific depletion from IgG EDDE (Figure 5A). Comparison to previously measured levels of PSMA+ EVs in the post-DRE samples showed some discrepancies from the MSD data, suggesting differences in PSMA protein vs. PSMA+ EV content of urine (sample 000-243-MDS).

**Figure 5.**
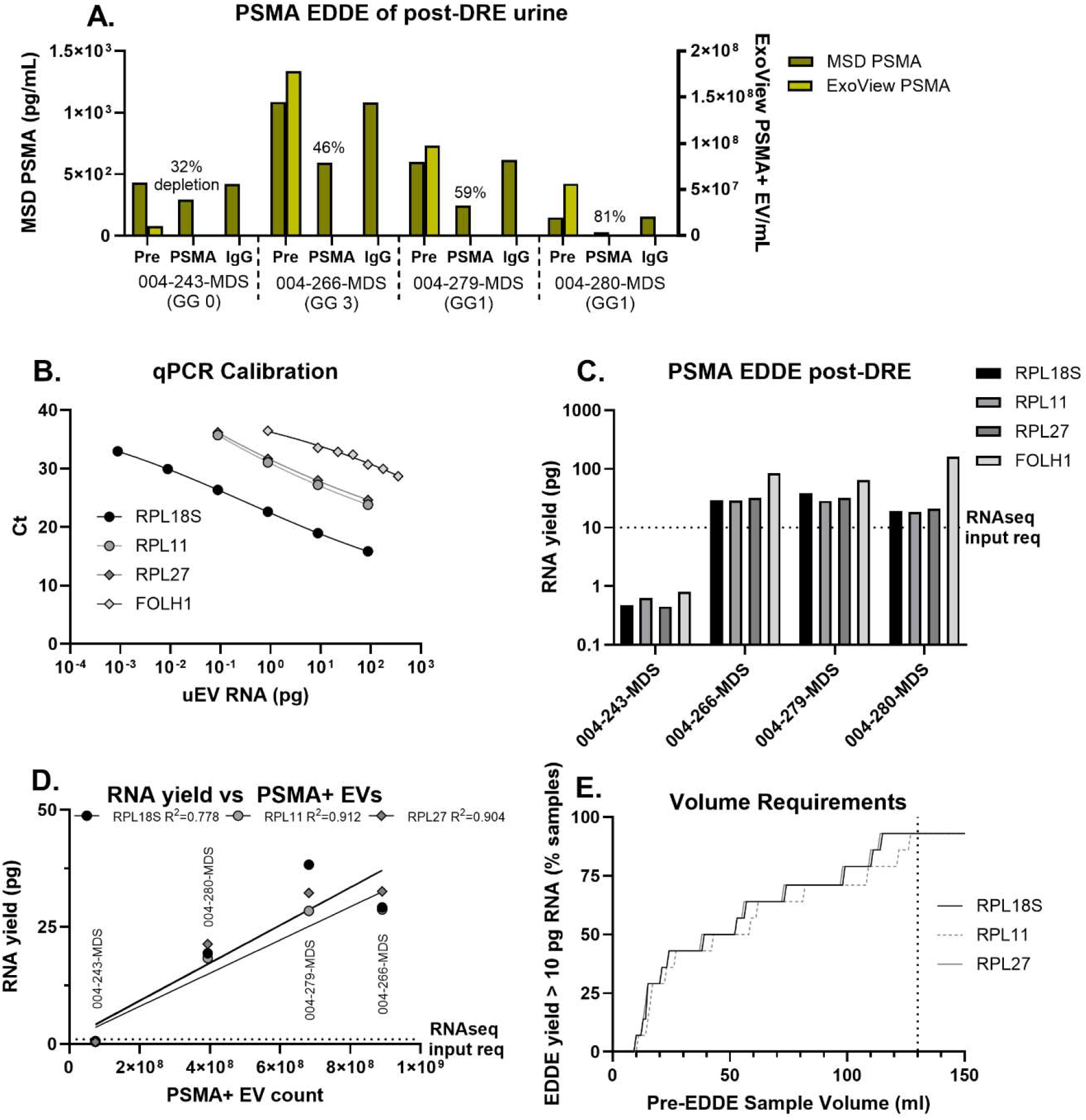
PSMA EDDE enriches sufficient RNA for sequencing from post-DRE urine. Panel A – PSMA levels measured in post-DRE samples via ExoView (light grey bars), or via MSD for the entire EDDE process (dark grey bars). Percent depletion of each sample is noted in the figure. Panel B – qPCR calibration curves used for RNA quantification. Serial dilutions of RNA extracted from urine EVs were measured with RPL18S, RPL11, RPL27 and FOLH1 qPCR assays, and picograms of RNA was measured on a BioAnalyzer for an undiluted stock solution. Panel C – Quantity of RNA enriched by PSMA EDDE from four post-DRE urine samples, quantified with qPCR. The minimum amount required as input for RNA sequencing using single-cell kit is shown as a dashed line. Panel D – Correlation of RNA yield from PSMA EDDE with the PSMA+ EV count in post-DRE urine. Fit quality (R^2^) is shown in the figure legend. Panel E – Saturation curves showing urine volume predicted to yield sufficient RNA from EDDE PSMA for sequencing, based upon PSMA+ EV concentration in pre-DRE (n=4) and GG0 (n=10) urine samples. Separate curves are shown for RPL18S, RPL11 and RPL27 PCR assays. The dotted line shows a cutoff of 130 mL, which is predicted to obtain > 10 pg/ml RNA for > 90% of the samples.

To enable accurate quantification of RNA yield, we generated qPCR calibration curves using serial dilutions of RNA extracted from urine EVs. Ct values for ribosomal genes RPL18S, RPL11, RPL27, and the prostate-specific gene FOLH1 (the coding gene of PSMA) were plotted against known RNA inputs, with RNA concentration of the undiluted stock determined by BioAnalyzer (Figure 5B). These curves were used to back-calculate RNA yield in experimental samples.

RNA extracted from PSMA-enriched EVs yielded detectable levels of all four transcripts across samples (Figure 5C). Importantly, RNA yield exceeded the minimum input requirement for single-cell RNA sequencing (dashed line), demonstrating the suitability of PSMA EDDE for downstream transcriptomic profiling.

To assess the relationship between EV abundance and RNA recovery, we plotted RNA yield against PSMA+ EV counts measured in pre-EDDE urine samples. A positive correlation was observed (Figure 5D), indicating that RNA yield scales with the number of immunocaptured EVs and supporting the utility of PSMA EDDE for isolating transcript-rich vesicles. We used the linear relationship between PSMA+ EV concentration in urine and RNA yield from PSMA EDDE to estimate the urine volume required to enable sequencing from non-DRE samples (Figure 5E). Using the measured PSMA+ EV counts in GG0 PCa urine (Figure 3B) and pre-DRE urine (Figure 3D), we can estimate that a volume of 130 mL should result in > 10 pg/ml of RNA for > 90% of the samples, suggesting that DRE is not necessary for RNA sequencing of PSMA+ EVs if larger urine volumes are collected.

## Discussion

This study establishes PSMA EDDE as a technically rigorous method for enriching prostate-derived EVs from urine, enabling transcriptomic analysis of tissue-specific EV populations. Immuno-isolation of EVs from complex biofluids remains a significant challenge due to low target abundance and the presence of confounding particles (Nieuwland & Siljander, 2024). To address this, we implemented a quantitative framework for tracking EV stoichiometry throughout the EDDE workflow, allowing precise measurement of depletion efficiency and yield. Our approach adheres to MISEV guidelines for EV characterization, including surface marker profiling and size distribution analysis via ExoView (Welsh et al., 2024). Given the low concentration of tissue-specific EVs in urine and the complexity of the matrix (Erdbrügger et al., 2021), such quantitative rigor is essential for optimizing immunocapture strategies and ensuring reproducibility across samples and conditions.

Through a systematic evaluation of the candidate surface markers PSMA, SLC45A3 and KLK2 present on EVs within PCa urine samples, we identified PSMA as the most abundant target for prostate EV enrichment. STEAP1 has been reported as another prostate-specific target used to profile prostate EVs in nanoflow cytometry (Kim et al., 2022), however we were unable to find reliable commercial antibodies for this target. We observed a significantly lower total EV count and lower PSMA+ EV count (which did not reach significance) in GG5 urine compared to GG0. Loss of ductal structure in prostate tissue in late disease could drive a decrease in total EV release into the urine, and we expect that middle grade groups (GG1 – GG4) would show an increase in EV concentration (Khoo et al., 2024). Total EV and PSMA+ EV abundance increased significantly following digital rectal exam (DRE), consistent with prior studies (Duijvesz et al., 2015; Khoo et al., 2024; Stephan & Jung, 2016). This supports the utility of post-DRE urine for prostate-specific EV isolation. However, this technique can be burdensome in the clinic. We found CD81+ EVs to be substantially lower concentration in urine than CD63 or CD9, a distinct pattern from that found in plasma (data not shown).

We have quantitatively analyzed the population rate of PSMA+ EVs within prostate cancer cell lines and urine from PCa donors. Using Exoview, we find that PSMA+ EVs comprise a very low percentage of the total EVs in LNCaP, ≤0.5%. We have also measured the amount of soluble PSMA protein in LNCaP EVs using the MSD assay and interestingly find a higher number of PSMA molecules per EV than expected based on ExoView measurements (Table 1). The discrepancy between the two platforms may be caused by soluble PSMA contained within purified LNCaP EVs, or PSMA internal to the lumen of EVs. ExoView claims to have nearly single molecule sensitivity, suggesting this is not the cause of the discrepancy. A third and most intriguing explanation is that EVs exist that contain PSMA, but not tetraspanins (which are necessary for detection on ExoView).

**Table 1.** Characterization of PSMA within LNCaP EVs.

|  |  |  |  |
| --- | --- | --- | --- |
| PSMA concentration (pg/mL) | 73970.8 |  |  |
| Leprechaun capture spot | CD63 | CD81 | CD9 |
| Total EV concentration (EV/mL) | $7.6 \times 10^{11}$ | $2.1 \times 10^{12}$ | $2.1 \times 10^{12}$ |
| PSMA+ EV concentration (EV/mL) | $2.7 \times 10^9$ | $8.2 \times 10^9$ | $1.1 \times 10^{10}$ |
| %PSMA+ EV | 0.4% | 0.4% | 0.5% |
| PSMA molecules per total EV<br>(assumed MW = 80kD) | 0.7 | 0.3 | 0.3 |

To establish the reliability of the EDDE workflow, we first validated positive and negative process controls using CD81- and IgG-directed EDDE on purified LNCaP EVs. CD81 EDDE consistently depleted >90% of CD63+, CD81+, and CD9+ EVs from the supernatant, suggesting a homogenous distribution of these markers in LNCaP-derived EVs. This was not the case in more complex biofluids such as plasma or urine, in which CD81 EDDE preferentially depleted CD81+ EVs (data not shown). The isotype control (IgG EDDE) showed minimal non-specific depletion (≤20%), confirming the specificity of the capture reagents. Eluate recovery of tetraspanin EVs was modest (≤25%), and not attributable to low pH-induced vesicle damage (Fig. 2E), suggesting that inefficient elution—rather than surface protein degradation—limits recovery. Notably, the downstream RNA workflow circumvents this limitation by lysing EVs on-bead, eliminating the need for elution and releasing captured materials directly for molecular analysis.

We evaluated four PSMA antibody clones—LNI-17, 460407, a polyclonal anti-PSMA, and 3/A12—by assessing their ability to bind EVs derived from PSMA-positive (LNCaP) and PSMA-negative (PC3) cell lines. Clones LNI-17 and 460407 exhibited the highest specificity and signal-to-noise ratio, making them strong candidates for EV enrichment. While several PSMA-targeting antibodies have been explored for prostate cancer diagnostics and therapeutics—such as 7E11 (used in the FDA-approved ProstaScint) and J591 (investigated for radiotherapy and EV detection via nanoflow cytometry (Britton et al., 2025; Davis et al., 2005; Tagawa et al., 2010) — these clones were not suitable for our application. Specifically, 7E11 binds the intracellular domain of PSMA, precluding its use for surface-based EV capture, and J591 was not commercially available. Nevertheless, clone 460407 demonstrated >90% depletion of PSMA+ EVs in EDDE experiments with LNCaP-derived vesicles, confirming its utility for effective prostate-specific EV isolation

Acidic elution conditions lead to substantial loss of PSMA+ EVs in the EDDE process, consistent with the description of PSMA on UniProt (Accession number Q04609) and prior reports of PSMA instability below pH 6.5 (The UniProt Consortium et al., 2025). To confirm that PSMA+ EVs were effectively captured and retained on the EDDE beads, we developed an on-bead immunoassay targeting PSMA. This assay verified successful retention of >80% of PSMA protein from the pre-enrichment sample, despite the absence of detectable PSMA in the eluate. Although intact EV recovery was compromised under these conditions, our RNA workflow does not rely on elution; instead, vesicles are lysed on-bead for direct access to cargo.

Application of PSMA EDDE to post-DRE urine samples showed sample dependent depletion efficiency ranging from 32 to 81%, as measured with the MSD-PSMA assay. We see a discrepancy between the concentration of PSMA protein (MSD assay) and PSMA+ EVs (ExoView) in the urine samples, as noted for LNCaP EVs (Table 1). We found a better correlation between PSMA+ EVs and RNA yields than soluble PSMA protein.

Using calibrated qPCR assays targeting three ribosomal genes and the prostate-specific transcript FOLH1, we quantified RNA yield from PSMA EDDE applied to post-DRE urine samples. Although overall yields were modest, three of the four samples exceeded 10 pg of RNA—meeting the input threshold for commercial sequencing platforms such as the SMART-Seq Total RNA-Seq Single Cell kit (Takara, catalog #634360). These results demonstrate the feasibility of downstream transcriptomic profiling from PSMA-enriched EVs. Moreover, the observed positive correlation between PSMA+ EV counts and RNA yield reinforces the quantitative reliability of the EDDE workflow. Modeling RNA recovery as a function of urine volume further suggests that sequencing from non-DRE samples is achievable with sufficient input, expanding the potential clinical utility of this approach.

Collectively, our findings position PSMA EDDE as a technically robust and clinically relevant strategy for non-invasive molecular profiling of prostate-derived EVs. Future studies will focus on sequencing RNA from both total urine EVs and PSMA-enriched subsets across clinical cohorts, enabling direct comparison of biomarker performance. This will help determine whether tissue-specific EV isolation offers meaningful advantages over bulk EV analysis. The ability to selectively enrich and interrogate prostate-derived EVs from urine opens new avenues for biomarker discovery, disease monitoring, and mechanistic studies in prostate cancer.

## Acknowledgements

This work was supported by funding from NCI / NIH R01CA272766.

## Supplemental Information

**Supplemental Figure 1.**
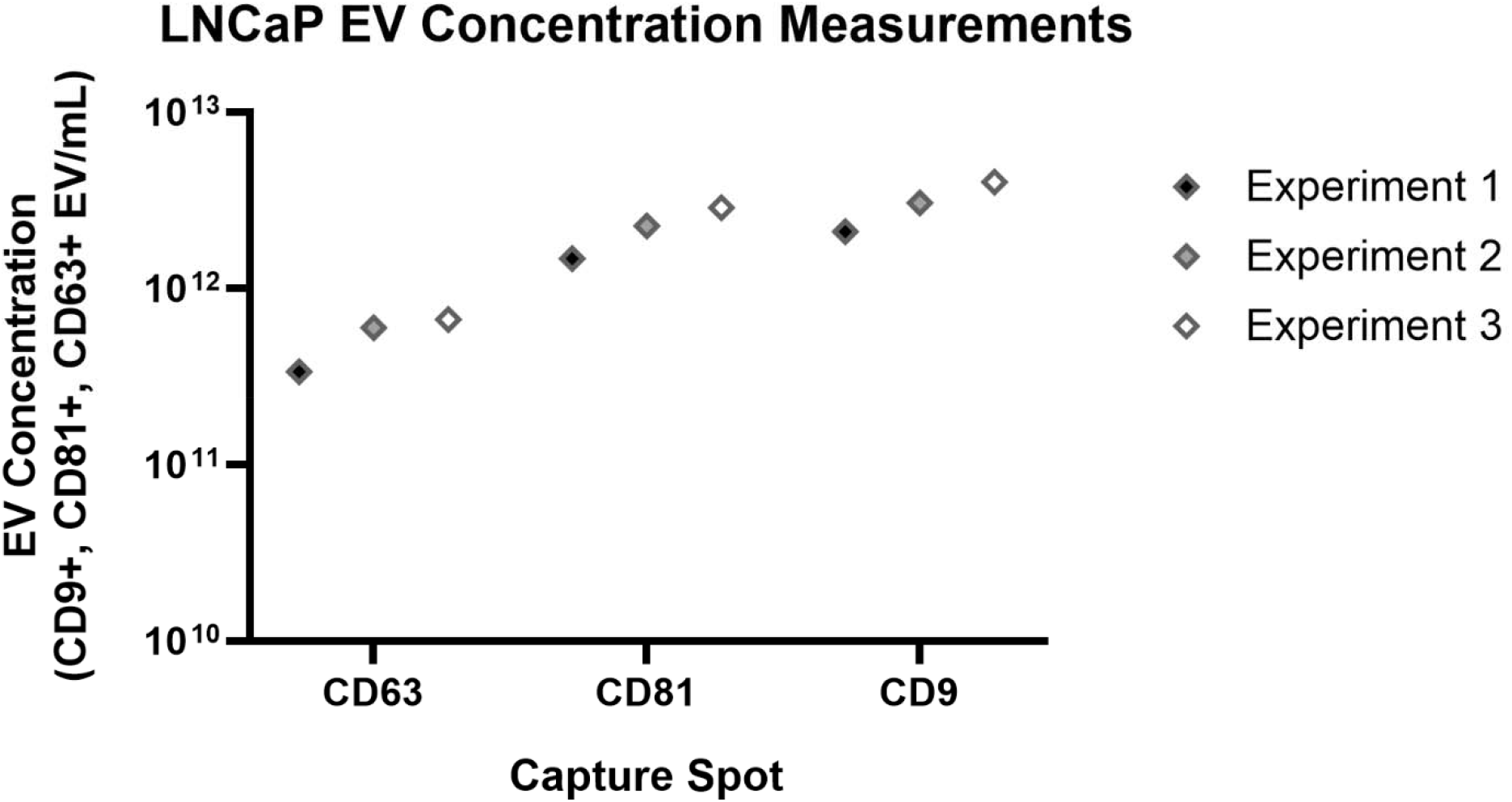
The average concentration of LNCaP EVs is approximately 1 × 10^12^ EV/mL. The concentration of LNCaP EVs was measured on ExoView in three independent experiments. A detection cocktail of CD63, CD81 and CD9 antibodies were used to measure tetraspanin+ EVs, reported as total EVs.

**Supplemental Figure 2.**
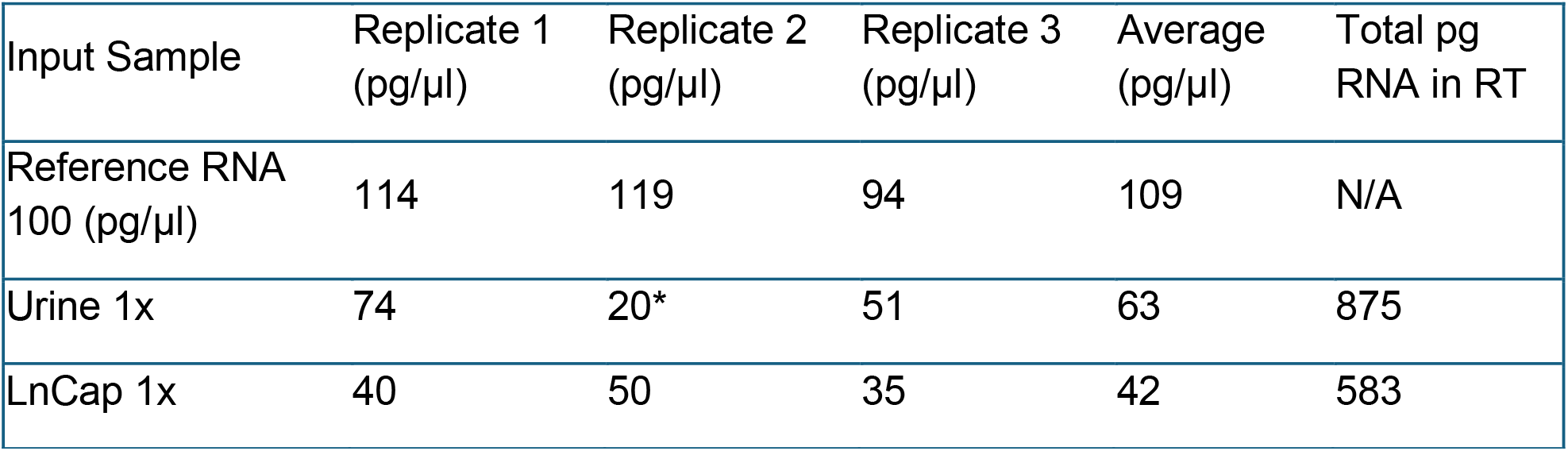
RNA quantition on BioAnalyzer. RNA was quantified from reference RNA (100 pg/µl), urine EVs (10 mL) and LnCap EVs (2 µl) in triplicates using the Bioanalyzer RNA Pico. Average of triplicates was used to determine input used in qPCR. For urine derived RNA, one replicate (marked with ^*^) was excluded due to being an outlier.

**Supplemental Figure 3.**
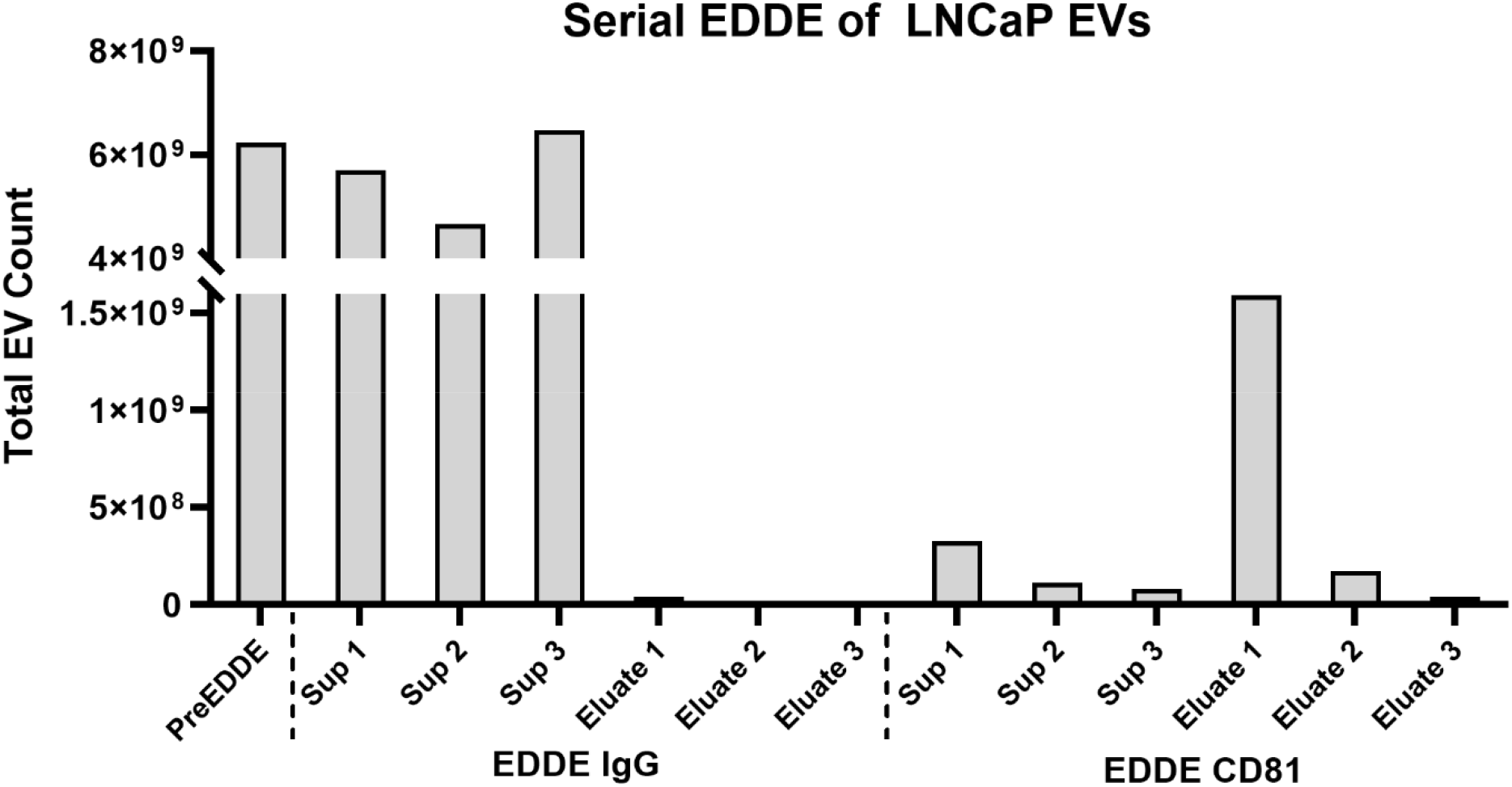
Serial EDDE captures LNCaP EVs in a sequentially increasing manner. Multiple rounds of EDDE CD81 and EDDE IgG (negative control) were performed on LNCaP EVs. EDDE CD81 sequentially depletes more of the supernatant, although the first round is sufficient to deplete >90% of EVs. Similarly, the eluate yield decreases sequentially with each round as well. Depletion and yield in eluate from IgG EDDE is insignificant, demonstrating the specificity of EDDE. Total EVs were measured with a cocktail of detection antibodies to CD63, CD81 and CD9 EVs, and counts were averaged across CD63, CD81 and CD9 capture probes.

**Supplemental Figure 4.**
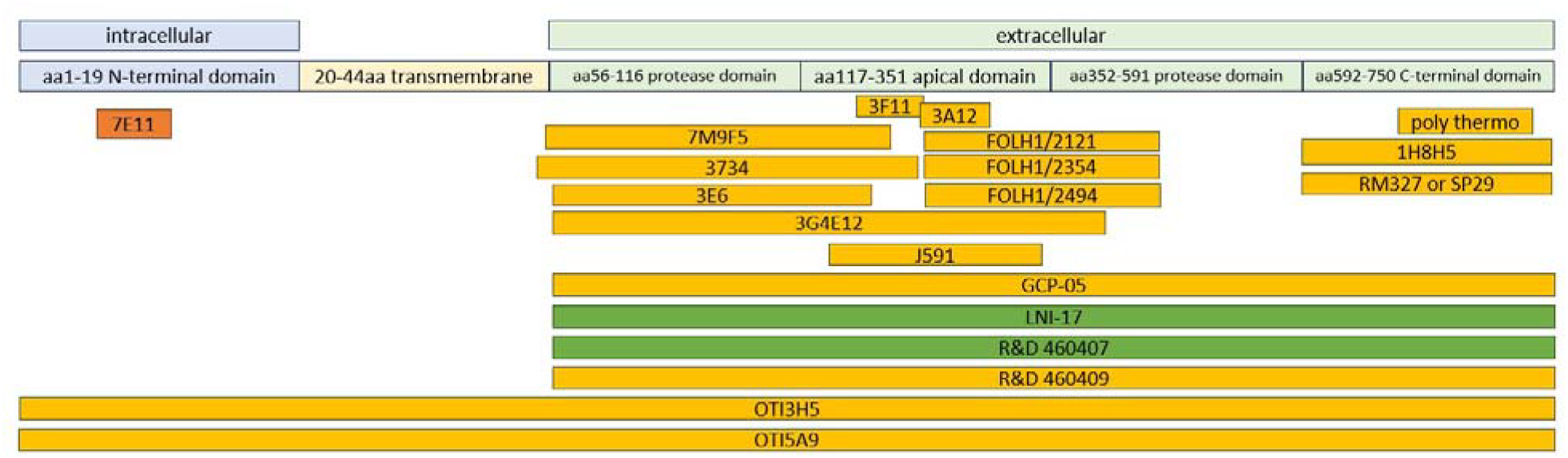
Epitopes of PSMA antibodies cover both the intra- and extra-cellular domains. Antibodies used for PSMA EDDE targeted only the extracellular domain. PSMA structural information was gathered from Uniprot, and antibody epitope information gathered from literature or vendor sources.

